# A Bayesian Approach for Estimating Branch-Specific Speciation and Extinction Rates

**DOI:** 10.1101/555805

**Authors:** Sebastian Höhna, William A. Freyman, Zachary Nolen, John P. Huelsenbeck, Michael R. May, Brian R. Moore

## Abstract

Species richness varies considerably among the tree of life which can only be explained by heterogeneous rates of diversification (speciation and extinction). Previous approaches use phylogenetic trees to estimate branch-specific diversification rates. However, all previous approaches disregard diversification-rate shifts on extinct lineages although 99% of species that ever existed are now extinct. Here we describe a lineage-specific birth-death-shift process where lineages, both extant and extinct, may have heterogeneous rates of diversification. To facilitate probability computation we discretize the base distribution on speciation and extinction rates into *k* rate categories. The fixed number of rate categories allows us to extend the theory of state-dependent speciation and extinction models (*e.g.,* BiSSE and MuSSE) to compute the probability of an observed phylogeny given the set of speciation and extinction rates. To estimate branch-specific diversification rates, we develop two independent and theoretically equivalent approaches: numerical integration with stochastic character mapping and data-augmentation with reversible-jump Markov chain Monte Carlo sampling. We validate the implementation of the two approaches in RevBayes using simulated data and an empirical example study of primates. In the empirical example, we show that estimates of the number of diversification-rate shifts are, unsurprisingly, very sensitive to the choice of prior distribution. Instead, branch-specific diversification rate estimates are less sensitive to the assumed prior distribution on the number of diversification-rate shifts and consistently infer an increased rate of diversification for Old World Monkeys. Additionally, we observe that as few as 10 diversification-rate categories are sufficient to approximate a continuous base distribution on diversification rates. In conclusion, our implementation of the lineage-specific birth-death-shift model in RevBayes provides biologists with a method to estimate branch-specific diversification rates under a mathematically consistent model.

## Introduction

Multiple lines of evidence unambiguously demonstrate that rates of diversification change over time and among lineages. The fossil record, for one, shows a pattern in which some groups flourish for a time, only to go extinct. Such a pattern cannot be explained by a constant-rate speciation and extinction model of cladogenesis (birth-death process). Once a group becomes reasonably speciose, it becomes almost impossible for it to die off unless the relative rates of speciation and extinction change. And, of course, the fossil record shows periods of time in which the rate of extinction dramatically increases for all lineages of the tree of life. But even without a fossil record, we would know that speciation and extinction rates have varied across the branches of the tree of life because the pattern of species richness in different groups differs so dramatically. How can the exceptional diversity of groups such as beetles or cichlids be explained except by an increased rate of diversification in those groups?

> *An inordinate fondness for beetles*
>
> *— J.B.S. Haldane, in Hutchinson (1959)*

Increasingly, questions regarding diversification-rate variation are pursued by inferring the parameters of explicit birth-death process models from phylogenies. For example, recent theoretical work has provided formal statistical phylogenetic methods that allow us to detect tree-wide changes in diversification rate, where the rates of all contemporaneous lineages vary either in a continuous manner (*e.g.,* Morlon et al. 2011; Etienne and Haegeman 2012; Condamine et al. 2013; Morlon 2014; Höhna 2014; Condamine et al. 2018), or in an episodic manner (*e.g.,* Stadler 2011), including episodes of mass extinction (*e.g.,* Höhna 2015; May et al. 2016). Similarly, formal statistical methods have been developed that allow us to infer state-dependent variation in diversification rates, where rates of lineage diversification are correlated with the state of a discrete character (*e.g.,* Maddison et al. 2007; FitzJohn 2012; Magnuson-Ford and Otto 2012; Beaulieu and O’Meara 2016; Freyman and Höhna 2018), or the value of a continuous trait (FitzJohn 2010).

In contrast to the methodological progress for studying tree-wide and state-dependent rate variation, efforts to develop methods for detecting variation in diversification rates across lineages have proven far more challenging. Rather than attempting to explicitly model shifts in diversification rates, early approaches for detecting among-lineage diversification-rate variation were based on summary statistics (Moore et al. 2004; Chan and Moore 2005) that do not provide estimates of branch-specific diversification rates. More recent approaches are motivated by birth-death processes using phylogenies (*e.g.,* MEDUSA by Alfaro et al. (2009) and BAMM by Rabosky (2014)) but contain mathematical errors (*i.e.,* the likelihood functions are incorrect). The reliability and robustness for parameter estimation of these methods is hotly debated (May and Moore 2016; Moore et al. 2016; Rabosky et al. 2017; Meyer and Wiens 2018; Meyer et al. 2018; Rabosky 2018; Barido-Sottani et al. 2018). The key problem is that none of the existing methods (Rabosky 2014; Barido-Sottani et al. 2018) take diversification-rate changes on extinct lineages into account. The omission of diversification-rate changes on extinct lineages is biologically problematic because: (a) extant species affected by a diversification-rate change might go extinct in the future and hence the diversification-rate change on a currently extant lineage might be a diversification-rate change on an extinct lineage in the future; and (b) the majority of species that ever existed (approximately 99%) has gone extinct which means that more diversification-rate changes must have occurred on extinct lineages. Even if we do not consider extinct lineages in our phylogenies, it is still crucial to model diversification-rate changes on extinct lineages because the probability of extinction fundamentally depends on the (changing) diversification rates in our models (Kendall 1948; Nee et al. 1994b,a).

Here, we develop a new Bayesian approach for inferring branch-specific rates of speciation and extinction. To this end, we first introduce the lineage-specific birth-death-shift process; a model that allows diversification rates to vary across the lineages of a phylogeny. Importantly, our lineage-specific birth-death-shift model rectifies the omission of diversification-rate changes on extinct lineages. We then extend previous theoretical work on inferring state-dependent diversification-rate variation to develop a numerical algorithm for computing the probability of the tree. We develop two theoretically equivalent approaches for estimating branch-specific rates of speciation and extinction; the first approach uses numerical integration together with stochastic character mapping and the second approach uses data augmentation together with reversible-jump Markov chain Monte Carlo sampling. All previous methods rely only on a data-augmentation approach which we show is less efficient. More importantly, we can validate our implementation and the underlying theory by demonstrating that estimates under the two equivalent approaches are, in fact, identical. Furthermore, we perform a simple simulation study which shows that our implementation behaves as one expects from Bayesian statistical theory. Finally, we explore the behavior of our method using an empirical example analysis of primates. All of the methods described in this paper have been implemented in the Bayesian phylogenetic inference software package RevBayes (Höhna et al. 2016).

## Methods

### The Lineage-Specific Birth-Death-Shift Process

We define a stochastic process that generates phylogenies via three events: (1) speciation events; (2) extinction events, and; (3) diversification-rate shift events. These events occur with rates *λ*_*i*_, *µ*_*i*_ and *η* respectively, where the index *i* stands for the *i*-th species. When a speciation event occurs, a lineage gives rise to two daughter lineages that inherit the speciation and extinction rates of their parent lineage. When an extinction event occurs, the lineage is simply terminated. When a diversification-rate shift occurs, new speciation and extinction rates are drawn from the corresponding base probability distributions, *f_*λ*_* (·) and *f_*µ*_*(·), and the lineage continues to diversify under these new rates. This defines a stochastic branching process in which rates of diversification are allowed to vary across lineages. We refer to this stochastic branching process as the lineage-specific birth-death-shift process.

Next, we explain how to simulate under the lineage-specific birth-death-shift process. This explanation has two purposes: (a) to clarify how the process works, and (b) to show that one can obtain realizations under the process which is sufficient to show that the process is in itself coherent. We imagine maintaining a list of ‘active’ lineages in computer memory. Under this stochastic branching process, the *i*-th active lineage can either speciate (with rate *λ*_*i*_) or go extinct (with rate *µ*_*i*_), and all active lineages can experience a diversification-rate shift (with a common rate *η*). We simulate the process over an interval, *T*, starting with one active lineage at time *t* = *T* in the past. The waiting times between events are exponentially distributed (because the probability of an event happening at a given time is equal if the rates are equal). Thus, we simulate forward in time by drawing an exponentially distributed waiting time for each active lineage. The parameter of the exponential distribution is the sum of the three event rates, (*λ*_*i*_ + *µ*_*i*_ + *η*). We pick the lineage with the shortest waiting time for the next event. We randomly choose the type of event for this lineage, which will be a speciation event with probability *λ*_*i*_/(*λ*_*i*_ + *µ*_*i*_ + *η*), or an extinction event with probability *µ*_*i*_/(*λ*_*i*_ + *µ*_*i*_ + *η*), or a diversification-rate shift event with probability *η*/(*λ*_*i*_ + *µ*_*i*_ + *η*).

When a lineage speciates, it is removed from the active list and replaced with its two daughter lineages, where each daughter lineage inherits the same speciation and extinction rates of their parent lineage. When a lineage experiences extinction, it is simply removed from the list of active lineages. When a diversification-rate shift occurs, the new speciation and extinction rates are drawn from the corresponding base probability distributions, *f_*λ*_* (·) and *f_*µ*_* (·), such that diversification rates are lineage specific. The simulation continues until the next event occurs after the present (*i.e., t ≤* 0), or until all lineages have gone extinct before time *t* = 0.

### Computing the Probability of an Observed Tree Under the Lineage-Specific Birth-Death-Shift Model

In outline, our method to compute the probability of an “observed” tree under the lineage-specific birth-death-shift model involves two components: (1) discretization of the speciation-and extinction-rate base probability distributions into *k* categories, to approximate the underlying continuous distributions; (2) a backwards algorithm that traverses the tree from the tips to the root in small time steps, ∆*t*. In each interval, we solve a pair of ordinary dffierential equations (ODEs) that compute the change in probability associated with all of the possible events (speciation, extinction, and diversification-rate shifts among the *k* diversification-rate categories) that could occur within each interval. Upon reaching the root, this algorithm has computed the probability of realizing the observed tree under each of the *k* discrete rate categories. Below, we detail each of these two components.

#### Discretization of the diversification-rate distributions

The probability calculations for the lineage-specific birth-death-shift model are impractical if we have to integrate over continuous base distributions for the diversification-rate parameters. Accordingly, we adopt an approach that provides an approximation of these integrals. Under this approach, we first divide the continuous probability distributions for the diversification-rate parameters into a finite number of *k* bins. The width of each bin (or diversification-rate category) is defined such that each category contains equal probability (*i.e.,* using the *k* quantiles of the underlying continuous probability distribution). Thus, the diversification rate for *i*-th discrete category is the median value of the corresponding quantile. As detailed in the following sections, our probability calculations involve summing over these *k* discrete diversification-rate categories.

As in the case of the discrete-gamma model for accommodating among-site variation in substitution rates (Yang 1994), the number of categories, *k*, is not a parameter of our model (*i.e.,* it is an assumption of the analysis rather than an estimate from the data). The choice of *k* categories represents a compromise: the resemblance to the underlying continuous probability distribution improves as the number of discrete categories increases (Figure 1). However, the computational burden also scales with the number of discrete categories. Thus, the value of *k* is only of interest to the extent that it must be sufficiently large to avoid discretization bias, while remaining small enough to allow practical computation. We will explore the impact of different numbers of diversification-rate categories in a later section.

**Figure 1:**
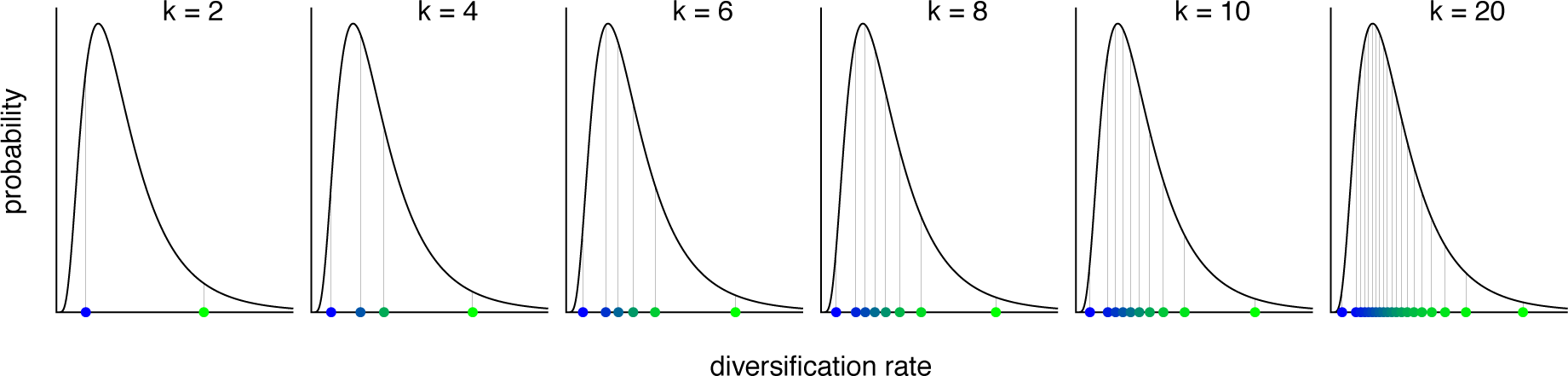
Approximation of the continuous base distributions for the diversification-rate parameters using discrete rate categories. Our approach for computing the probability of the data under the lineage-specific birth-death-shift model specifies *k* quantiles of the continuous base distributions for the speciation and extinction rates. We compute probabilities by marginalizing (averaging) over the *k* discrete rate categories, where the diversification rate for a given category is the median of the corresponding quantile (colored dots). This approach provides an efficient alternative to computing the continuous integral, and will provide a reliable approximation of the continuous integral when the number of categories *k* is sufficiently large to resemble the underlying continuous distribution.

#### Backwards algorithm to compute the probability of the observed phylogeny

The second part of our approach involves discretizing the tree into tiny time steps, and then numerically integrating over these time slices to compute the probability of the observed data under the lineage-specific birth-death-shift process. This aspect of our approach draws heavily on the algorithm developed by Maddison et al. (2007) and FitzJohn (2012) in the context of exploring a state-dependent birth-death process (their BiSSE and MuSSE model). Following Maddison et al. (2007), our numerical algorithm begins at the tips of the tree where *t* = 0 (*i.e.,* the present). We need to consider two probability terms at each point in time: *D*(*t*) and *E*(*t*). *D*(*t*) is the probability of the observed lineage between time *t* and the present, and *E*(*t*) is the probability that a lineage at time *t* goes extinct before the present. For each tip, we must initialize *D*(*t*) and *E*(*t*) and also consider the state of the process. Under the BiSSE model, the diversification process depends on the state of the binary character (0 or 1). Thus, for a species with the observed state 0, we initialize *D*_0_(0) = 1 and *D*_1_(0) = 0. Conversely, for a species with the observed state 1, we initialize *D*_0_(0) = 0 and *D*_1_(0) = 1. Under our model, the state of the diversification process is not observed. Thus, for each species at time *t* = 0, we initialize *D_i_*(0) = 1 for each of the *i ∈* (1,…, *k*) discrete diversification-rate categories. In fact, this is equivalent to the case under the BiSSE model when the state of a given species is unknown (*i.e.,* coded as ‘?’), in which case we would initialize *D*_0_(0) = 1 and *D*_1_(0) = 1. Finally, we initialize the extinction probability for each species as *E_i_*(0) = 0 for each of the *i ∈* (1,…, *k*) discrete diversification-rate categories. Note that if we have an incomplete (but random/uniform) sample of species, then we would initialize *D_i_*(0) = *ρ* and *E_i_*(0) = 1 − *ρ* for each of the *i ∈* (1,…, *k*), where *ρ* is the proportion of randomly sampled species (FitzJohn et al. 2009).

Next, we begin our traversal of the tree from each tip (where *t* = 0) to the root in tiny time steps, ∆*t*. For each time step into the past, we calculate the change in probability of the observed lineage over the interval (*t*+∆*t*) by enumerating all of the events that could occur within the interval ∆*t*. If we assume that ∆*t* is small, then the probability of any two events occurring in the same interval is negligible. In the interval ∆*t* there are four possible scenarios that could occur (see Equation 1 and Figure 2): (*i*) nothing happens (no speciation event or diversification-rate shift), or (*ii*) no diversification-rate shift but a speciation event occurs and the left descendant subsequently goes extinct before the present, or (*iii*) no diversification-rate shift but a speciation event occurs and the right descendant subsequently goes extinct before the present, or (*iv*) no speciation event occurs but there is a diversification-rate shift to any of the other (*k −*1) rate categories. Now we can compute *D_i_*(*t*+∆*t*) by writing the set of *k* difference equations *D*_1_(*t*+∆*t*)*, D*_2_(*t*+∆*t*)*,…, D_k_*(*t*+∆*t*):

**Figure 2:**
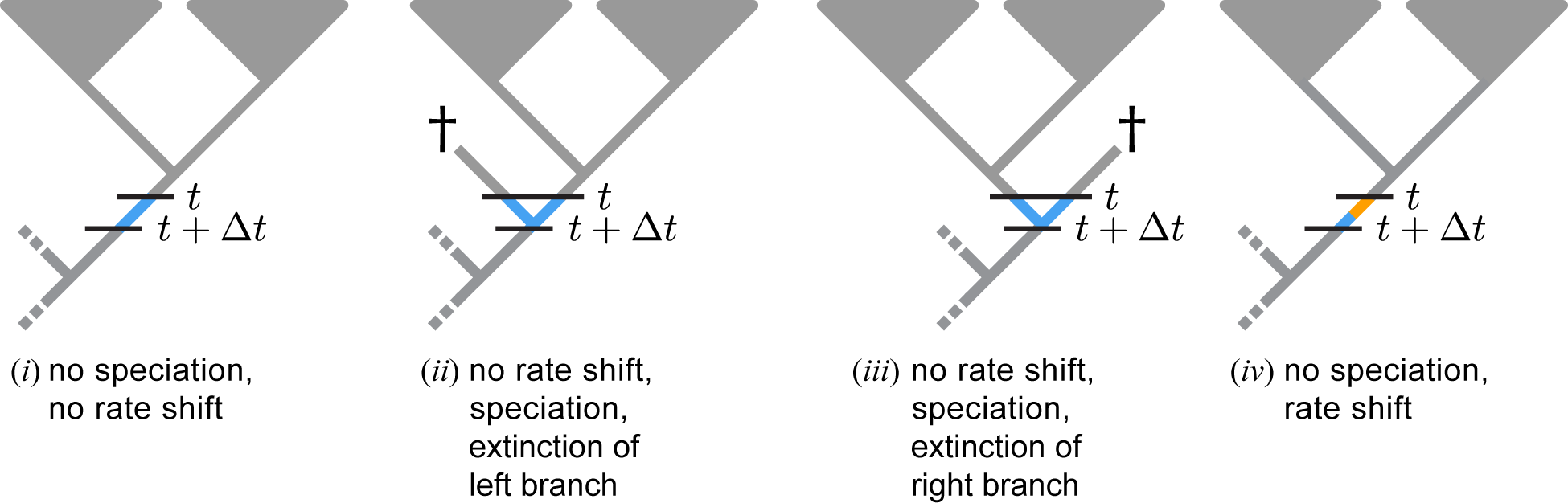
Possible scenarios that could occur over the interval ∆*t* along a lineage that is observed at time *t*. To compute the probability under the lineage-specific birth-death-shift process, we traverse the tree from the tips to the root in small time steps, ∆*t*. For each step into the past, from time *t* to time (*t* + ∆*t*), we compute the change in probability of the observed lineage by enumerating all of the possible scenarios that could occur over the interval ∆*t*: (*i*) nothing happens, (*ii*) a speciation event occurs, where the right descendant survives and the left descendant goes extinct before the present, or (*iii*) a speciation event occurs, where the left descendant survives but the right goes extinct before the present, or (*iv*) a diversification-rate shift from category *i* to *j* occurs. Color key: segment(s) of the tree within the interval ∆*t* are colored blue for state *i* and/or orange for state *j* to reflect the conditioning of the corresponding scenarios, segment(s) of the tree between *t* and the present are colored gray because we have integrated over the *k* discrete rate categories (no specific assignment of rate categories), and segments of the tree between *t*+∆*t* and the root are colored gray because we will integrated over the *k* discrete rate categories.

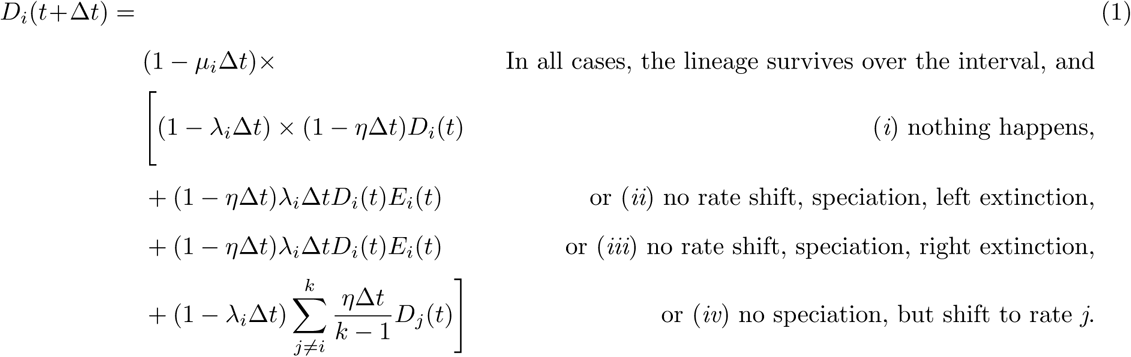

Note that the first (unnumbered) term in Equation 1 represents the probability that the observed lineage does not go extinct in the interval ∆*t*. The probability of no extinction in the interval ∆*t* is included because if the lineage had gone extinct in this interval, then we could not have observed it.

Equation 1 makes it clear that in order to compute *D_i_*(*t*), we must simultaneously compute *E_i_*(*t*) (the probability of a lineage going extinct before the present). Again, we calculate the change in the extinction probability for each step into the past, from *t* to (*t* + ∆*t*), by enumerating all of the scenarios that could occur within the interval ∆*t* (see Equation 2 and Figure 3): (*i*) in the first scenario, the lineage goes extinct in the interval, ∆*t*; in the remaining scenarios, the lineage does not go extinct in the interval, and (*ii*) the lineage does not speciate and does not experience a diversification-rate shift during the interval ∆*t*, but subsequently goes extinct before the present, which occurs with probability *E_i_*(*t*), or (*iii*) the lineage speciates in the interval, ∆*t*, such that both descendent lineages must eventually go extinct before the present, which occurs with probability *E_i_*(*t*)^2^, or (*iv*) the lineage does not speciate in the interval, ∆*t*, but does experience a diversification-rate shift from category *i* to category *j*, and subsequently goes extinct before the present, which occurs with probability *E_j_*(*t*). As before, we can compute *E_i_*(*t*+∆*t*) by writing the set of *k* difference equations *E*_1_(*t*+∆*t*)*, E*_2_(*t*+∆*t*)*,…, E_k_*(*t*+∆*t*):

**Figure 3:**
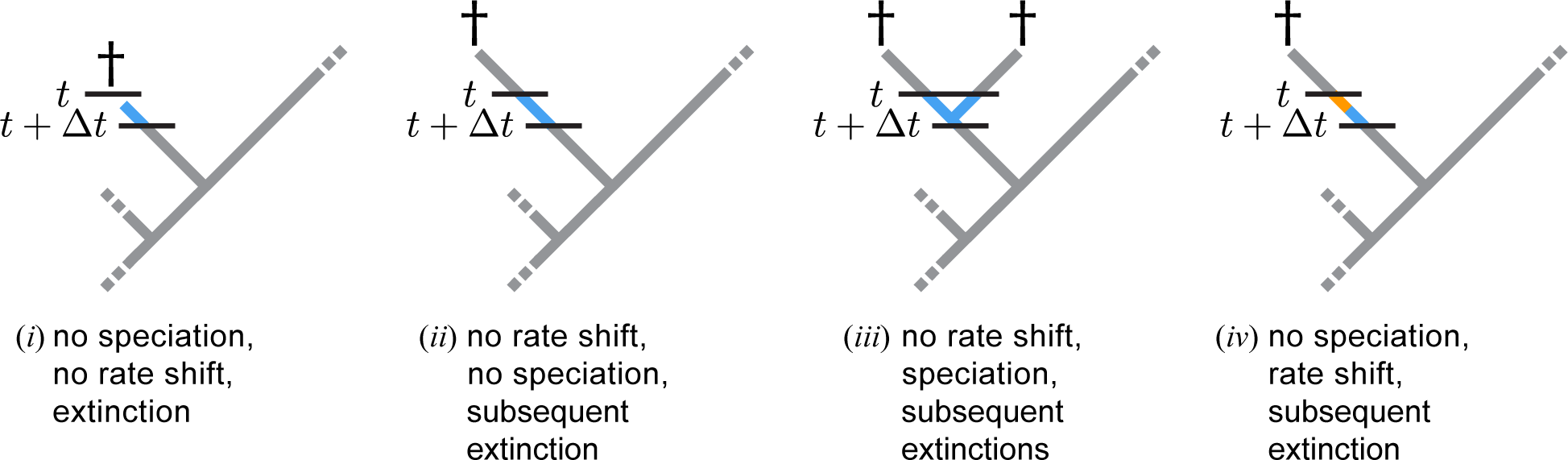
Possible extinction scenarios. For each step into the past, from time *t* to time (*t*+∆*t*), we compute the change in the extinction probability, *E*_i_(*t*) (the probability that a lineage in state *i* at time *t* goes extinct before the present) by enumerating the scenarios that could occur in the interval ∆*t*: (*i*) the lineage goes extinct in the interval ∆*t*; in the remaining three scenarios, the lineage does not go extinct in the interval, and (*ii*) nothing happens (no extinction, speciation or diversification-rate shift in the interval ∆*t*), with subsequent extinction before the present, (*iii*) the lineage speciates in the interval ∆*t*, with subsequent extinction of both daughter lineages before the present, or (*iv*) the lineage experiences a diversification-rate shift from rate category *i* to *j*, with subsequent extinction before the present. Segments of the tree are colored as described in the key for Figure 2.

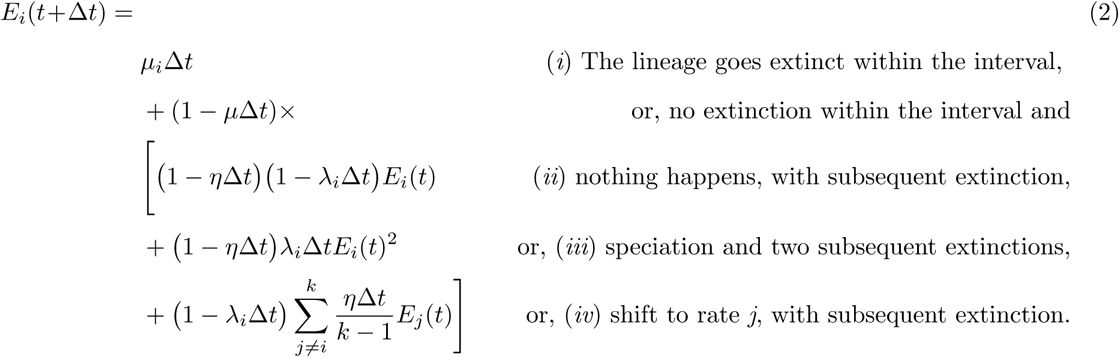

We now derive the ordinary differential equations from the corresponding difference Equations 1 and 2. This requires some algebra (which includes dividing by the interval ∆*t* and omitting terms of order (∆*t*)^2^) and results in the coupled ordinary differential equations (ODEs):

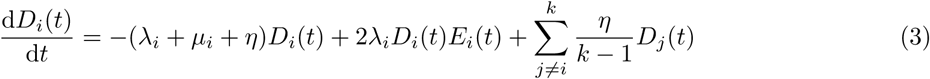

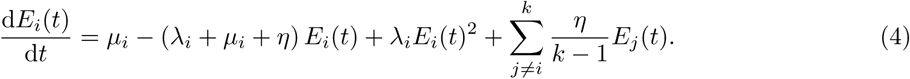

These differential equations are solved for each branch of the phylogeny and compute the probability of an observed lineage. As an aside, we note that we store the values of *D_i_*(*t*) and *E_i_*(*t*) computed at some interval, ∆*δ*. We will use these stored values for the procedure that maps diversification-rate shifts over the tree (see the description of the forwards algorithm, below).

Because we are moving backward in time, each branch will end at the speciation event by which it originated. For a speciation event that occurs at time *t* while the process is in diversification-rate category *i*, we initialize the probability density of the immediately ancestral lineage, *A*, by taking the product of its two daughter species at time *t* 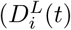 and 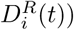 multiplied by the probability density of the observed speciation event at time *t*, *λ*_*i*_:

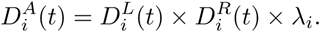

The algorithm terminates when we reach the most ancient speciation event in the tree (*i.e.,* at the root). Upon reaching the root of the tree, we will have computed the vector of *k* probabilities, *D_i_*(*T*), where *i ∈* {1, 2,…, *k*}. *D_i_*(*T*) is the probability of observing the entire tree under the lineage-specific birth-death-shift process given that the process was initiated in diversification-rate category *i* at the root. We then multiply each of these *k* probabilities by their corresponding prior probabilities, *π*_*i*_. The prior probability for rate category *i* specifies the probability that the diversification process started in category *i* at the root. Recall that each of the *k* discrete diversification-rate categories has equal probability (*i.e.,* they are quantiles of the corresponding base distributions). Therefore, we assume that all of the *k* diversification-rate categories have equal prior probability, *π*_*i*_ = 1*/k* (*i.e.,* a discrete uniform prior distribution). The product of the root probability for diversification-rate category *i* and the prior probability for diversification-rate category *i* gives the probability of rate category *i*. Finally, the sum of these *k* probabilities gives the probability of the entire tree under the lineage-specific birth-death-shift model

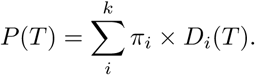

We will call this probability *P* (*T*) of the ‘observed’ phylogeny the likelihood function under the numerical integration approach because we perform parameter estimation in a Bayesian statistical framework.

### Estimating Branch-Specific Speciation and Extinction Rates using Stochastic Character Mapping (forward algorithm)

The backwards algorithm computes the probability of the observed tree under the lineage-specific birth-death-shift process. In doing so, however, the numerical marginalization ‘integrates out’ the focal parameters: the branch-specific diversification rates. Therefore, we adopt an approach to estimate the branch-specific rates of speciation and extinction that is based on stochastic character mapping (Huelsenbeck et al. 2001; Nielsen 2002; Landis et al. 2018; Freyman and Höhna 2019). Under stochastic character mapping, character histories are simulated in a forwards traversal of the tree (*i.e.,* moving over the tree from the root to its tips), where each history specifies the number, location and magnitude of character-state changes. Here, we adopt the algorithm developed by Freyman and Höhna (2019) for mapping diversification histories. The objective is to compute the probability that the diversification process is in each of the *k* rate categories, *F_i_*(*t−∆t*). To compute *F_i_*(*t−∆t*) we take the product of three probability components: the initial probabilities of the *i* rate categories at the beginning of the interval, *F_i_*(*t*), the forward probabilities of the process over the interval *∆t*, and the conditional likelihoods of the process between (*t−∆t*) and the present, *D*(*t−∆t*).

Our algorithm starts at the root of the tree, where we initialize the diversification process by randomly drawing one of the *k* rate categories proportional to their corresponding probabilities at the root, *P_i_*(*T*). Next, we initialize the forward probability *F_i_*(*t*) of the selected rate category with probability 1, and the other (*k −*1) rate categories have zero probability (*i.e., F_i_*(*T*) = 1 and *F_j≠i_*(*T*) = 0). Then, we begin our traversal in tiny time steps, ∆*t*, forward in time from time *t* to time (*t−∆t*). We calculate the probability *F_i_*(*t −∆t*) that the diversification process is in rate category *i* at time (*t −∆t*) by enumerating all of the scenarios that could occur within the interval ∆*t* that result in the lineage being in rate category *i* at time (*t−∆t*), given the initial state, *F_i_*(*t*) (see Equation 5). We have the same four scenarios as in Figure 2 and Equation 1, so we omit a repetition of the details here. The main difference is the direction of time (*i.e.,* we move forwards in time) and that the surviving lineage at time (*t−∆t*) must evolve into the lineage observed at the present, which occurs with probability *D_i_*(*t −∆t*). We compute *F_i_*(*t −∆t*) by writing the set of *k* difference equations *F*_1_(*t−∆t*)*, F*_2_(*t−∆t*)*,…, F_k_*(*t−∆t*):

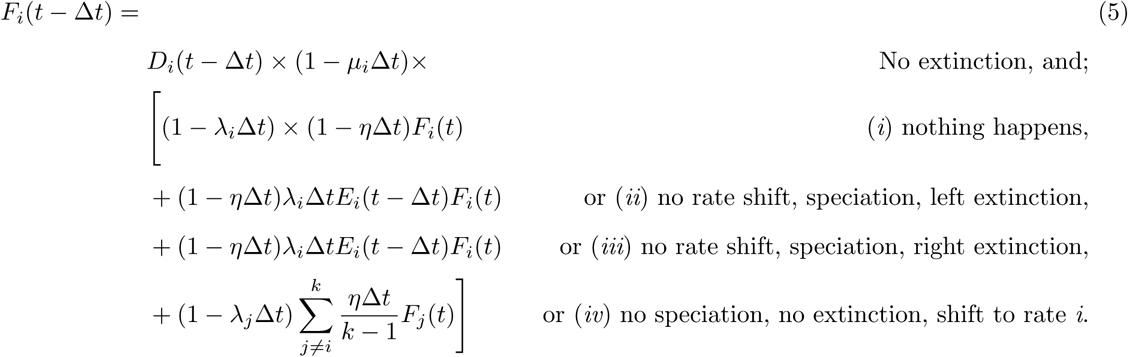

As previously, we derive the ordinary differential equation from its corresponding difference Equation 5 by using some algebra and omitting terms of order (∆*t*)^2^:

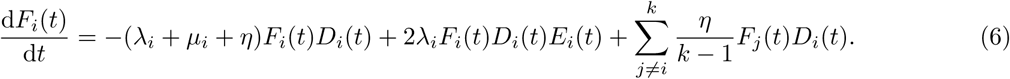

We compute these probabilities by solving this ODE in a forwards traversal of the tree. Specifically, at a given branch at time *t* where we just mapped the state *i*, we solve *F_i_*(*t*) until time (*t −* ∆*δ*). Note that ∆*t* is much smaller than ∆*δ* (∆*t ≪* ∆*δ*) because we take the limit of ∆*t →* 0 in the numerical integration but draw character maps only after a time step of ∆*δ*. Then, at time *t −∆δ*, we draw one of the *k* diversification-rate categories proportional to their corresponding probabilities, *F_i_*(*t −* ∆*δ*). The sampled rate category becomes *F_i_*(*t −* ∆*δ*) = 1 for the next iteration of the recursive forwards algorithm. If the rate category sampled at time (*t−∆δ*) is the same as the initial rate category (at time *t*), we paint the interval ∆*δ* of the branch by the corresponding diversification-rate category. Conversely, if the rate category sampled at time (*t−∆δ*) differs from the initial rate category (at time *t*), we paint a diversification-rate shift between these two rate categories within the interval ∆*δ*. The recursive algorithm continues moving forward in time and terminates upon reaching the tips of the tree. Upon reaching the present, we will have mapped a complete diversification-rate history that specifies the number and location of diversification-rate shifts and the rate category for each branch of the tree.

### An Alternative Approach Using Data Augmentation

Next, we develop a second numerical algorithm for estimating branch-specific diversification rates. Specifically, our second approach is based on data augmentation (Dempster et al. 1977; Tanner and Wong 1987; Gelfand and Smith 1990; Huelsenbeck et al. 2000; Landis et al. 2013; Uyeda and Harmon 2014), where we augment the study tree (*i.e.,* our actual data) with diversification histories (describing the number and location of diversification-rate shifts and the rate category for every branch of the tree). We treat these diversification histories as observations (*i.e.,* they augment our data). We compute the likelihood of each ‘observed’ diversification history using a modified version of our backwards algorithm. We then use reversible-jump MCMC (RJ-MCMC) to sample diversification histories in proportion to their posterior probability (see Appendix A).

Consider a tree that has been augmented with a history that specifies the diversification-rate category for every branch of the tree. As previously, we compute the probability of the observations (the phylogeny and the ‘observed’ diversification history) using a backwards algorithm that moves over the tree from the tips to the root in tiny time steps, ∆*t*. For each interval, we compute the probability of the data by solving a pair of ODEs that account for all of the scenarios that could occur over each step into the past. We begin at the tips of the tree, where *t* = 0 (the present), where we initialize the two probability terms, *D*(*t*) and *E_i_*(*t*). Observe that we use only a single probability term *D*(*t*) because a lineage that is in state *i* always has probability *D_j_*(*t*) = 0 for all other diversification rate categories *j*. For all species we initialize *D*(0) = 1 or in the case of incomplete sampling we initialize *D*(0) = *ρ*. Finally, we initialize the extinction probability for each species as *E_i_*(0) = 0 for each of the *i ∈* (1,…, *k*) diversification-rate categories (or in the case of incomplete sampling we initialize *E_i_*(0) = 1 − *ρ*).

Next, we calculate the probability of the observed lineage and the ‘observed’ diversification history over the interval (*t*+∆*t*) by enumerating all possible scenarios that could occur within the interval ∆*t*. When a diversification-rate shift is not ‘observed’ within the current interval, there are three possible scenarios that could occur over the interval (see Equation 7 and Figure 4A), specifically: (*i*) no speciation event occurs (*i.e.,* nothing happens), or (*ii*) a speciation event occurs and the left descendant subsequently goes extinct before the present, or (*iii*) a speciation event occurs and the right descendant subsequently goes extinct before the present. Accordingly, we can compute *D*(*t*+∆*t*) as a difference equation:

**Figure 4:**
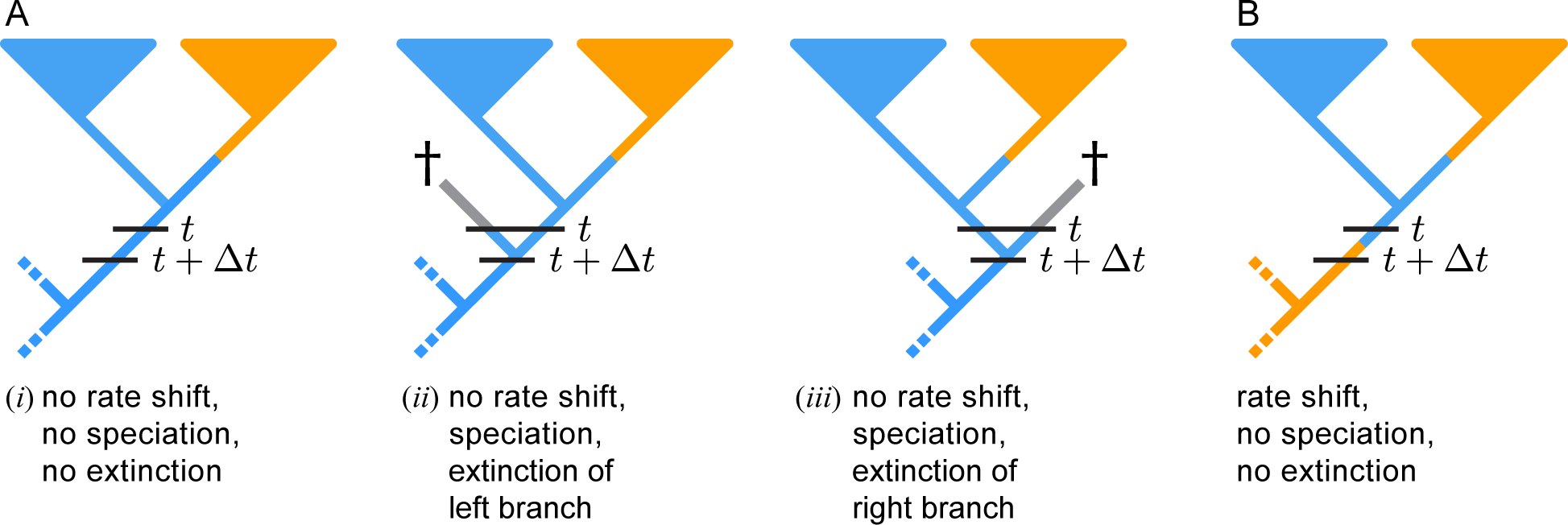
Possible scenarios that could occur over an interval ∆*t* under the data-augmentation approach. The observed phylogeny has been augmented with a diversification history (describing the number and location of rate shifts and the discrete rate category for every branch segment of the tree), which we treat as an observation. To compute the probability of the observed tree and the ‘observed’ history under the lineage-specific birth-death-shift process, we traverse the tree from the tips to the root in small time steps, ∆*t*. For each step into the past, from time *t* to time (*t*+∆*t*), we compute the probability of the observations by enumerating all of the possible scenarios that could occur over the interval ∆*t*. (A) When no diversification-rate shift is ‘observed’ in the interval ∆*t*, there are three scenarios: (*i*) nothing happens, or (*ii*) a speciation event occurs, where the right descendant survives and the left descendant goes extinct before the present, or (*iii*) a speciation event occurs, where the left descendant survives but the right goes extinct before the present. (B) Alternatively, a diversification-rate shift from category *i* to *j* is ‘observed’ within the interval ∆*t*. Color key: segments of extant lineages are colored according to the ‘observed’ diversification history (blue segments are in rate category *i*, orange segments are in rate category *j*); segments of the tree between *t* and an extinction event are colored gray because we average the extinction probabilities over the *k* discrete diversification-rate categories.

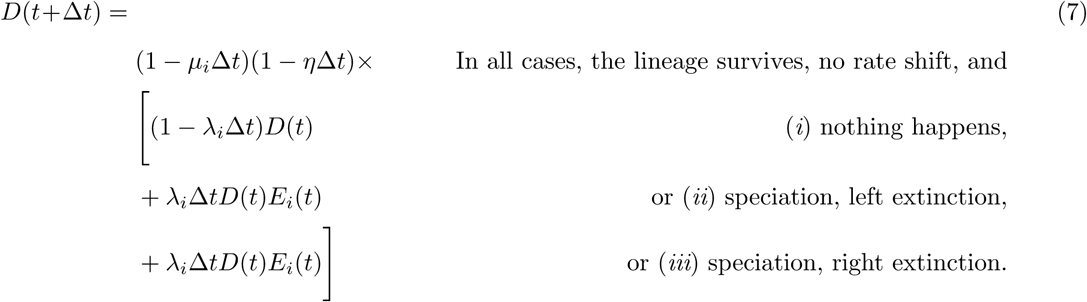

The first two (unnumbered) terms in Equation 7 account for the probability that the observed lineage does not go extinct in the interval ∆*t* (otherwise it could not have been observed at the more recent time, *t*), and also for the probability that the lineage does not experience a diversification-rate shift in the interval ∆*t* (because no diversification-rate shift was ‘observed’). Diversification-rate histories cannot be mapped onto unobserved (extinct) branches. Therefore, we compute extinction probabilities, *E_i_*(*t*), in exactly the same way as before (see Equations 2 and 4 and Figure 3).

As previously, we derive the ordinary differential equation from its corresponding difference Equation 7:

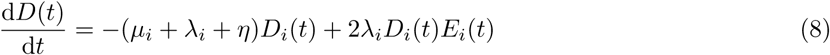

As previously, we compute the probability of the observations by solving these ODEs (*i.e.,* by integrating the change in probability over each time step, ∆*t*, from the present to time *t*).

We continue traversing the current branch toward the root of the tree (moving in small time steps, ∆*t*, further into the past, and solving the coupled ODEs for each interval) until we either reach the end of the branch (at a speciation event, in which case the probabilities are propagated as described previously), or we encounter a diversification-rate shift. When we encounter an ‘observed’ diversification-rate shift from category *i* to category *j* (where *i ∕*= *j*), we initialize *D′*(*t*) as:

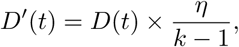

which is the current probability of the observed lineage multiplied by the probability density of ‘observing’ a diversification-rate shift to one of the other (*k −* 1) rate categories at time *t* (Figure 4B). The algorithm terminates when we reach the root of the tree. Since we are only considering one term *D*(*t*) for the observed lineages in any state *i*, this probability *D*(*t*) gives us directly the probability of observing the tree and diversification rate history. We will call this probability of the ‘observed’ phylogeny augmented with diversification histories the likelihood function under the data-augmentation approach because we perform parameter estimation in a Bayesian statistical framework.

## Validating the Theory and Implementation

We performed several tests to evaluate both the underlying theory and the implementation of the lineage-specific birth-death-shift model in RevBayes, including: (1) comparing analytical likelihoods to those estimated using the two methods under the special case where there are no diversification-rate shifts, (2) comparing analytical and empirical distributions of the number of diversification-rate shifts under the special case where all rate categories are identical, (3) comparing parameter estimates under the two theoretically equivalent but independent approaches, (4) assessing the computational efficiency of the two approaches, and (5) assessing the ability of the method to recover true parameter values under simulation. We briefly describe each of these experiments below (we provide further details of these analyses in the Supplementary Material and the scripts available online from https://github.com/hoehna/birth-death-shift-analyses).

### Comparing Analytical and Numerically Approximated Probabilities for the Special Case of a Constant-Rate Birth-Death Process

Recall that there is no analytical solution for computing the likelihood under the lineage-specific birth-death-shift process, which motivates the development of our two numerical algorithms. However, the likelihood can be computed analytically for the special case when *η* = 0 (*i.e.,* when the process simplifies to a constant-rate birth-death process). Thus, we compare the analytical likelihood to that approximated using the two numerical methods under the special case of a constant-rate birth-death process. If our derivation and implementation are correct, and we chose a sufficiently small ∆*t*, then the likelihoods should be exactly identical under the three different methods.

For the computations, we set all of the *k* diversification-rate categories equal, assumed *k* = 4 discrete rate categories, and set *η* = 0 (the rate of diversification-rate shifts). We then computed the likelihood over a range of relative-extinction rates, *ε* = {0*,…,* 1} using the analytical solution under the constant-rate birth-death process, the numerical-integration and data-augmentation methods. As expected, plots of the analytical and numerically approximated likelihoods are identical (Figure 5), confirming both the derivation and implementation of the two numerical algorithms.

**Figure 5:**
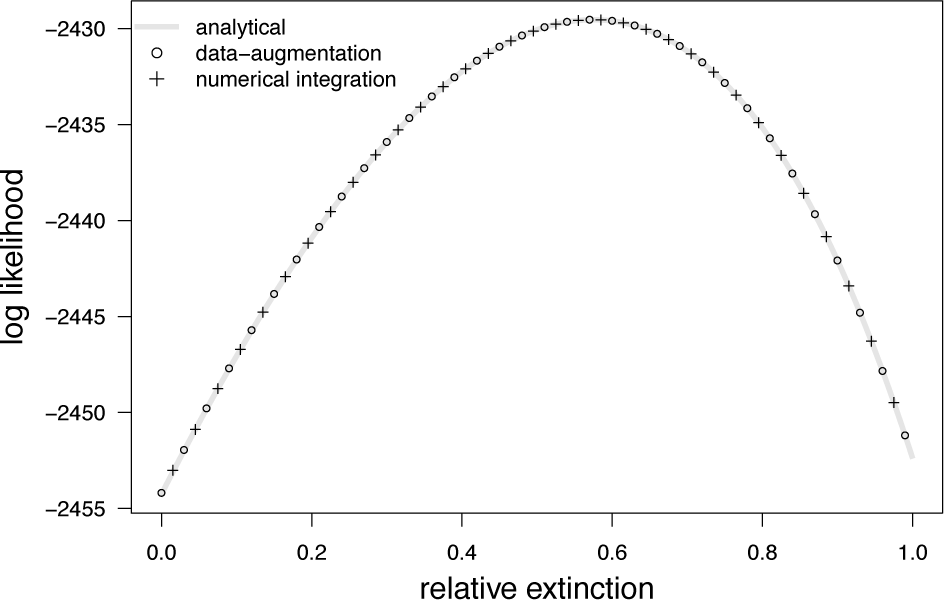
Comparing the analytical likelihoods to those approximated using the numerical algorithms when *η* = 0. We can analytically compute the likelihood under the special case where the rate of diversification-rate shifts is zero. We plot the analytical likelihood over a range of values for the relative-extinction rate, *ϵ* = *µ ÷ λ* (shaded line), and compare these values to those estimated using the numerical-integration method (symbols) and the data-augmentation method (+ symbols). The analytical and estimated likelihoods are identical, confirming the correctness of the derivation and implementation of the independent methods.

### Comparing Analytical and Estimated Distributions for the Number of Diversification-Rate Shifts

Second, we compare the analytical and estimated probability distributions on the number of diversification-rate shifts. Under the lineage-specific birth-death-shift process, waiting times between diversification-rate shifts are exponentially distributed with rate *η*. If we constrain the *k* diversification-rate categories to be equal, then diversification-rate shifts among those *k* identical rate categories will have no impact on the probability of speciation or extinction. The difference in the probability of the observed phylogeny stems only from the probability of the number of diversification-rate shift events but not the probability of speciation and extinction. In this case, the number of diversification-rate shifts over the branches of the tree is Poisson distributed with rate *η* × *TL* where *TL* is the tree length (*i.e.,* the sum of all of branch lengths in the tree).

We first plot the analytical distribution for the number of diversification-rate shifts over a set of values for the shift-rate prior that specify a corresponding range of values for the expected number of diversification-rate shifts, *E*(*S*) = {1, 10, 20}. Next, we estimate the posterior number of diversification-rate shifts using our two independent implementations. The distribution for the number of diversification-rate shifts estimated using either approach should follow the corresponding analytical distribution. As expected, plots of the analytical and estimated probability distributions for the number of diversification-rate shifts are identical (Figure 6), confirming that both numerical algorithms are correctly implemented in RevBayes. Moreover, this result does not only confirm our implementation of the probability of an observed phylogeny under the lineage-specific birth-death-shift model but specifically validates the MCMC algorithms to sample from the number of diversification-rate shift events under the prior distribution.

**Figure 6:**
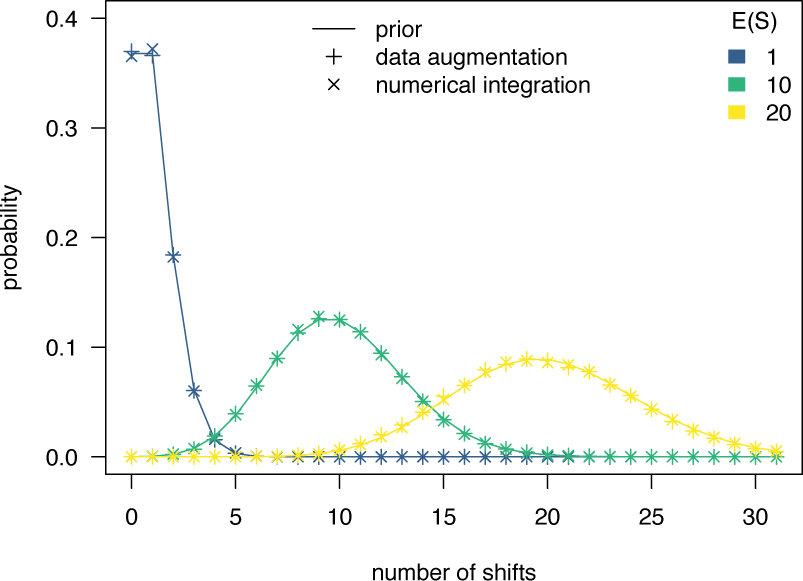
Distribution of the number of diversification-rate shifts when all categories have an identical diversification rate. The plot depicts the analytical distribution of the number of diversification-rate shifts over a set of values for the shift-rate, *η*, that specify a corresponding range of values for the expected number of diversification-rate shifts *E*(*S*) = {1, 10, 20}. We estimated the number of diversification-rate shifts using both the numerical-integration method (×symbols) and the data-augmentation method (+ symbols) for the same range of shift-rate priors when the diversification rate was specified to be the same for all of the *k* diversification-rate categories. The analytical and estimated distributions are identical, confirming the correctness of the derivation and implementation of the independent methods.

### Comparing Branch-Specific Parameter Estimates Between the Two Implementations

The data-augmentation and stochastic character mapping method for estimating branch-specific speciation and extinction rates rely on different likelihood functions as well as different MCMC algorithms. Nevertheless, both methods should provide the same estimated posterior distribution of branch-specific speciation and extinction rates. Therefore, we estimated branch-specific speciation and extinction rates using both methods and compared the results over a range of values for the number of discrete diversification-rate categories, *k* = {4, 6, 8, 10, 20}. The models for both analyses were set to be exactly the same so that we expected that branch-specific diversification rates are the same (up to some stochasticity due to the MCMC sampling procedure).

Figure 7 shows the estimated posterior mean of the branch-specific mean speciation rates. The estimates of the two alternative methods are nicely correlated. This correlation demonstrates that our derivation of the theory and implementation are (mostly likely) correct. It would have been very unlikely that we introduced the same mistake in the two independent methods giving the exact same bias. Note that this validation is stronger than comparing two independent implementations of the same method because we show that two different methods using different derivations of the likelihood yield the same results if applied to the same model.

**Figure 7:**
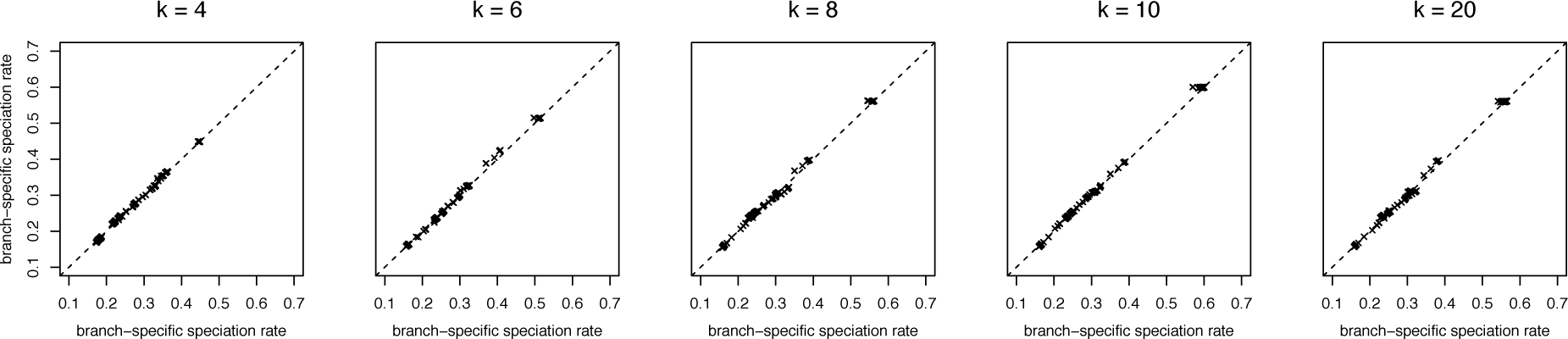
Comparison between branch-specific speciation rate estimates using data-augmentation and stochastic character mapping. We estimated branch-specific speciation and extinction rates using our data-augmentation and stochastic character mapping methods with *k* = {4, 6, 8, 10, 20} rate categories respectively. For each branch, we calculated the average speciation and extinction rates, *i.e.,* if there was a rate-shift event, then we computed the weighted average of the rates weighted by the time spent in a rate category. This plot shows the mean posterior estimates for both methods. As we expect, both method provide the same rate estimates.

### Computational Eciency of Data-Augmentation and Stochastic Character Mapping

The theory and derivation predicts that the data-augmentation and stochastic character mapping methods yield identical estimates of branch-specific diversification rates. We have established in Figure 7 that indeed both methods provide identical branch-specific diversification rate estimates. Until now, all implementations of similar methods use only a data-augmentation approach (Rabosky 2014; Barido-Sottani et al. 2018).

Since both approaches give identical estimates, we are interested in which method is computationally more ecient. We performed a set of MCMC analyses under identical model settings for both methods over a range of datasets (providing a range of tree sizes). We assessed the impact of (a) number of diversification-rate categories *k*, (b) the expected number of diversification-rate shifts *E*(*S*), and (c) the tree size.

The stochastic character mapping method outperforms the data-augmentation method with respect to higher effective sample size per CPU second (Figure 8). The main advantage of the stochastic character mapping method is that it does not need additional parameters such as the number, location/timing and magnitude of the diversification-rate shifts. Instead, the rate-shift events are directly sampled from the conditional posterior distribution, which is extremely efficient. It is therefore not surprising that the stochastic character mapping method is computationally superior. Indeed, we had considerable problems to obtain convergence using the data-augmentation method. Thus, we recommend biologists who are interested in estimating branch-specific diversification rates to use the stochastic character mapping method only and we will do so for the following sections.

**Figure 8:**
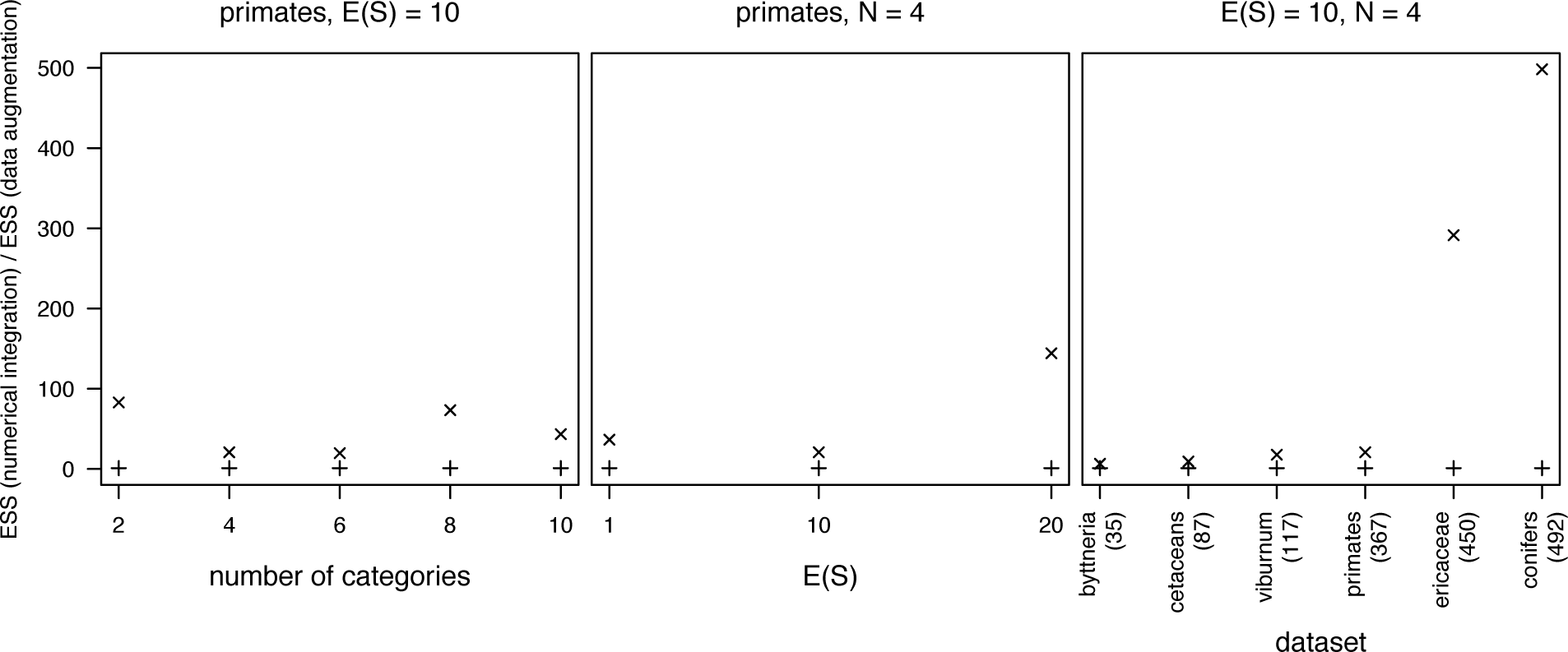
Comparison of MCMC performance between data augmentation and marginalization. We computed branch-specific diversification rates using our two implementations for the primates phylogeny for different number of rate categories (left) and different number of expected shift events (middle). Additionally, we used several different phylogenies to asses the impact of tree size (right). We plot here the effective sample size (ESS) of the numerical integration method normalized by the ESS of the data-augmentation method. Thus, we show the performance gain in MCMC efficiency of the numerical integration method compared to the data-augmentation method.

### Validation using Simulation

Our implementation of the lineage-specific birth-death-shift process in RevBayes allows for performing parameter inference and simulating under the process. Here we describe a small simulation study focused on confirming that our implementation is correct, and we leave exploring the model’s full range of statistical behavior under various diversification scenarios to future work. To this end, we simulated trees under the lineage-specific birth-death-shift process, estimated the branch-specific net-diversification rates using MCMC sampling, and confirmed that the credible intervals of our branch-specific net-diversification rates had the correct coverage.

We simulated 1000 trees under the lineage-specific birth-death-shift process using 4 rate categories conditional on having 200 surviving tips. We rather arbitrarily chose 200 surviving tips because these simulated datasets were not too small for reliable inference and yet still small enough to run reasonably fast. Trees were simulated in forward time until 201 lineages were alive. The trees were then trimmed back in time randomly within the interval between where there were 200 and 201 lineages. We then estimated the branch-specific diversification rates for each simulated tree using the numerical-integration method (more details about the simulation and inference settings are given in the Supplementary Material).

If our implementation of the lineage-specific birth-death-shift process and MCMC sampling machinery is implemented correctly, then we should obtain coverage probabilities that are equal to the width of the credible interval (Huelsenbeck and Rannala 2004). Here we used coverage probabilities as the proportion of times across the 1000 simulation replicates the credible interval of estimated branch-specific net-diversification rate contained the true simulated value. Figure 9 shows that coverage probabilities are equal to their corresponding credible intervals. Thus, we obtained more evidence that our software implementation is correct.

**Figure 9:**
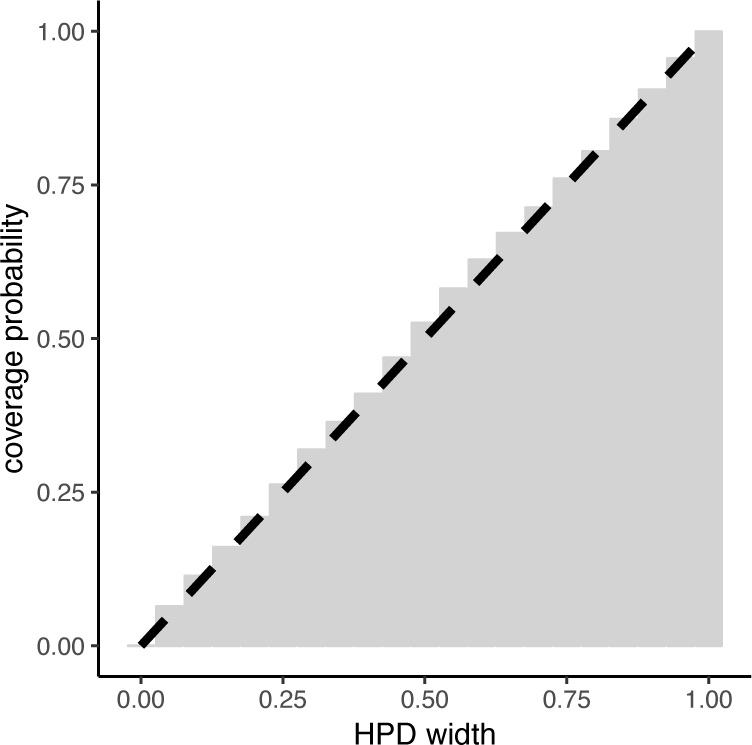
Coverage probabilities of branch-specific net diversification rate estimates for different credible interval widths. The coverage probabilities (*y*-axis) of branch-specific net-diversification rate estimates are plotted at different highest posterior density interval widths (*x*-axis). The coverage probabilities were calculated as the proportion of times across the 100 simulation replicates the credible interval contained the true simulated branch-specific net-diversification rate. If our model and the inference machinery is implemented correctly this should correspond with the diagonal line where *y* = *x* (dashed line).

Figure 10 illustrates one example of the simulation replicates used. This example demonstrates that the overall precision of estimated net-diversification rates is high. The method particularly has power to detect the location of diversification rate shifts when they lead to large clades. The method has little power to detect those diversification rate shifts that lead to small clades.

**Figure 10:**
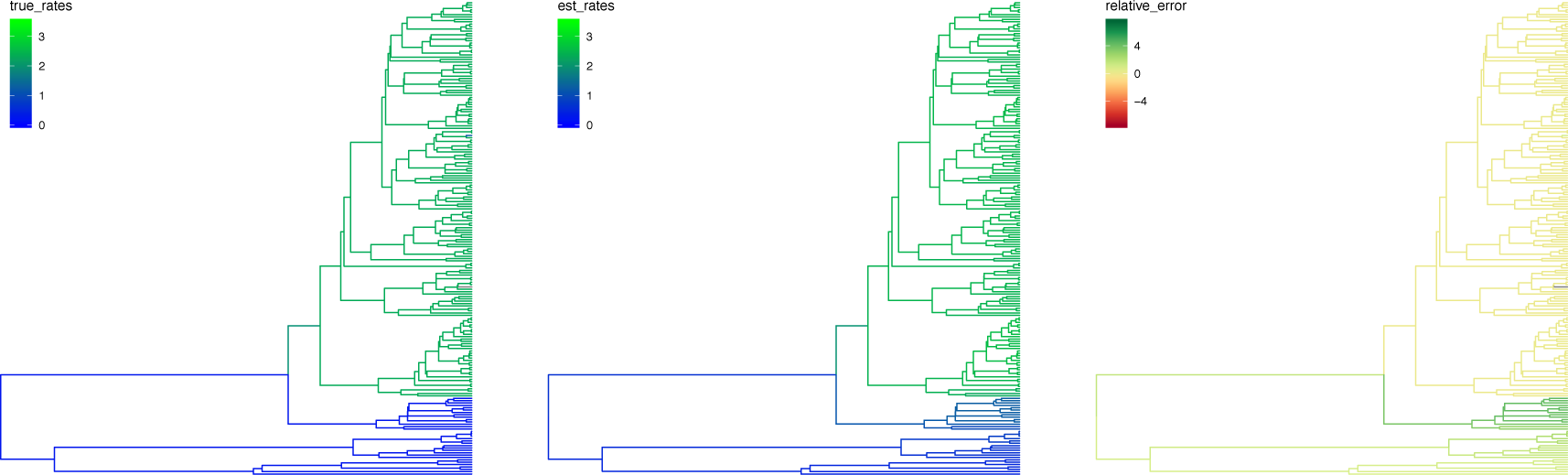
An example replicate from the simulation study. *Left:* A tree simulated using RevBayes under the lineage-specific birth-death-shift process with the branches colored to show the true mean branch-specific net diversification rates. *Center:* Estimates of the branch-specific net diversification rates made by RevBayes. Diversification rate shifts in large clades are accurately estimated, however diversification rate shifts in lineages leading to small clades were not detected due to the small number of branches resulting in a lack of power. *Right:* The precision of net diversification rate estimates measured as the relative error in the branch-specific rate estimates. The relative error is low throughout the tree except for places in which rate shifts occurred in small clades.

## Empirical Example Analysis of Primates

Next, we complement our method-validation with an exemplary analyses of an empirical primate phylogeny obtained from Springer et al. (2012). Our objective is to explore several important aspects of the lineage-specific birth-death-shift model, including: (1) assessing the sensitivity of branch-specific diversification-rate estimates to the assumed number of diversification-rate categories *k*, (2) assessing the sensitivity of posterior estimates of the number of diversification-rate shifts to the choice of shift-rate prior, and (2) assessing the sensitivity of posterior estimates of the branch-specific diversification rates to the choice of shift-rate prior. We briefly describe each of these experiments below (again, we provide further details of these analyses in the Supplementary Material and scripts available online).

### Robustness of Branch-Specific Diversification Rate Estimates to the Number of Diversification-Rate Categories

Recall that we approximate the continuous base distribution of the speciation and and extinction rate using discretization (Figure 1). The quality of this approximation depends on the chosen number of discrete rate categories. When we use a small number of categories, the estimates of the branch-specific speciation rates may be biased, but as the number of rate categories increases to infinity, the discretized process should converge toward the continuous one. Unfortunately, increasing the number of rate categories comes with some cost, as the time it takes to compute the probability of a tree is proportional to the number of rate categories.

Here we explored the impact of the number of diversification rate categories on branch-specific diversification rate estimates. Specifically, we esimated the branch-specific speciation rates for different numbers of rate categories, *k* = {2, 4, 6, 8, 10, 20}. Then, we compared the branch-specific speciation rate estimates of adjacent numbers of diversification rate categories (*i.e.,* 2 vs. 4, 4 vs. 6, etc.). Indeed, when the number of rate categories is low, branch-specific rate estimates are sensitive to the chosen number of rate categories (Figure 11, left panels). Encouragingly, as the number of rate categories increases, the branch-specific rate estimates converge toward the same values (Figure 11, right panels). These results suggest that an adequate approximation of the continuous distribution can be achieved with few diversification rate categories. In our case, 6 diversification rate categories seem to be a sufficient approximation but we choose 10 rate categories to be slightly conservative. As a general rule, using a *k* = 10 runs reasonably efficient while large values of *k* (*e.g.,* 100 or more) become computationally infeasable.

**Figure 11:**
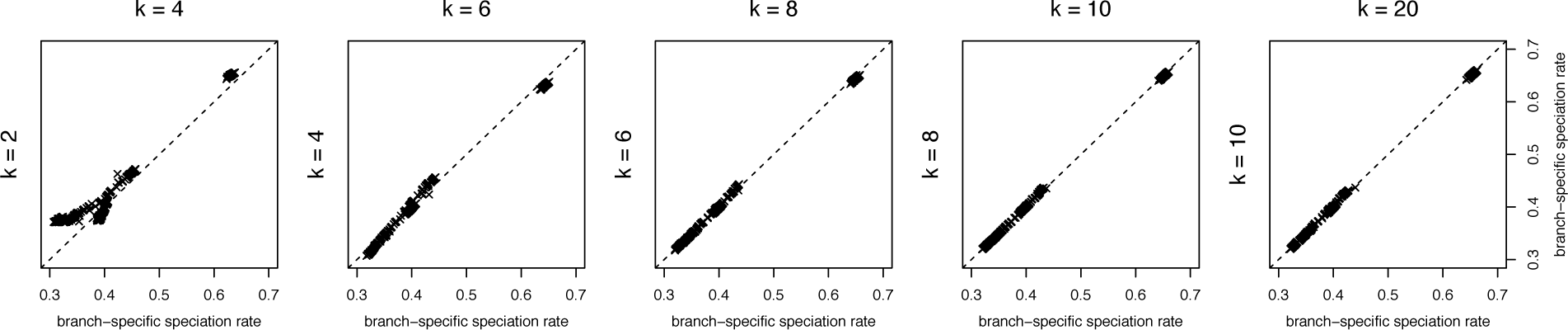
Comparison of branch-specific rate estimates for different numbers of diversification-rate categories. We estimated the posterior mean branch-specific speciation rate for each branch of the primate tree where the number of rate categories was set to *k* = {2, 4, 6, 8, 10, 20}. We then compared the mean estimates of the rates between adjacent pairs of the number of diversification-rate categories. For small numbers of diversification-rate categories, the branch-rate estimates are quite different between adjacent settings. However, as the number of categories increases, the branch-specific diversification-rate estimates converge toward stable estimates.

### Prior Sensitivity of the Estimated Number of Diversification-Rate Shifts

Previous work has shown that the inferred number of diversification-rate shifts in birth-death-shift models can be extremely sensitive to the prior on the rate of shifts (Moore et al. 2016). Therefore, we analyzed the primate phylogeny under a range of priors on *η* specified so that the expected number of diversification-rate shift events under a Poisson process was *E*(*S*) = {1, 10, 20}. For each shift-rate prior, we estimated the corresponding marginal posterior distribution for the number of diversification-rate shifts.

While the posterior number of diversification-rate shifts (slightly) departed from their respective prior distributions, they nevertheless are (very) sensitive to the prior (Figure 12). This results implies that estimates of the number of rate-shift events have to be treated carefully and are only meaningful in the context of their corresponding prior distribution. More work is needed to evaluate how robust estimates of the number of rate-shift events are and how much power there is to detect such events. In the meantime, we strongly recommend that researchers perform inference under a range of prior choices for the expected number of rate-shift events.

**Figure 12:**
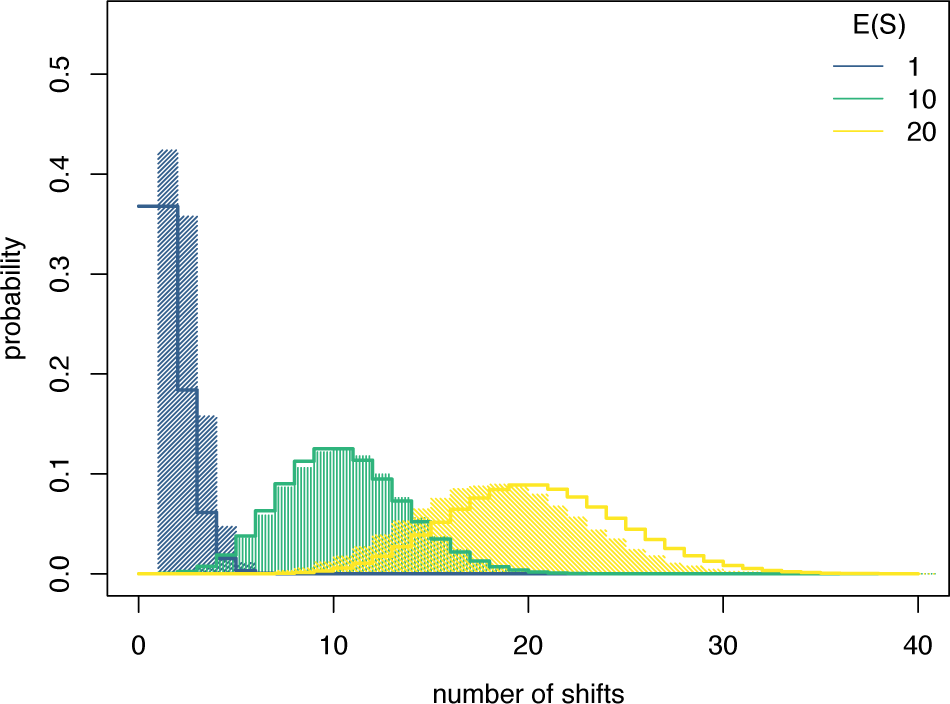
Comparison between the prior number of diversification-rate shifts and the posterior number of diversification-rate shifts for different shift-rate priors. We estimated the posterior number of diversification-rate shifts (shaded bars) in the primate phylogeny under three different shift-rate priors, with the prior on *η* specified so that the prior expected number of shifts under a Poisson process, *E*(*S*), was 1, 10, or 20 (solid lines). The posterior number of diversification-rate shifts is very sensitive to these prior settings although not exactly matching the prior distributions.

### Robustness of Branch-Specific Diversification-Rate Estimates to the Prior on the Expected Number of Diversification-Rate Shifts

Posterior estimates of the number of diversification-rate shifts are quite sensitive to the choice of shift-rate prior (Figure 12). However, it remains unclear whether other parameters (*e.g.,* branch-specific speciation rates) may also be similarly sensitive to the choice of shift-rate prior. To understand the robustness of branch-specific speciation-rate estimates to the prior on *η*, we compared the posterior means of branch-specific average speciation-rate parameters estimated under different prior values of *E*(*S*).

In contrast to the estimated number of diversification-rate shifts, the branch-specific diversification rate estimates are less sensitive to the prior on *η* (Figure 13). For example, in all cases we infer increased speciation rates in a subclade of the Old World Monkeys (Figure 14). We therefore recommend that biologists focus on the branch-specific diversification rate estimates as the the parameter of interest because we can estimate them more robustly.

**Figure 13:**
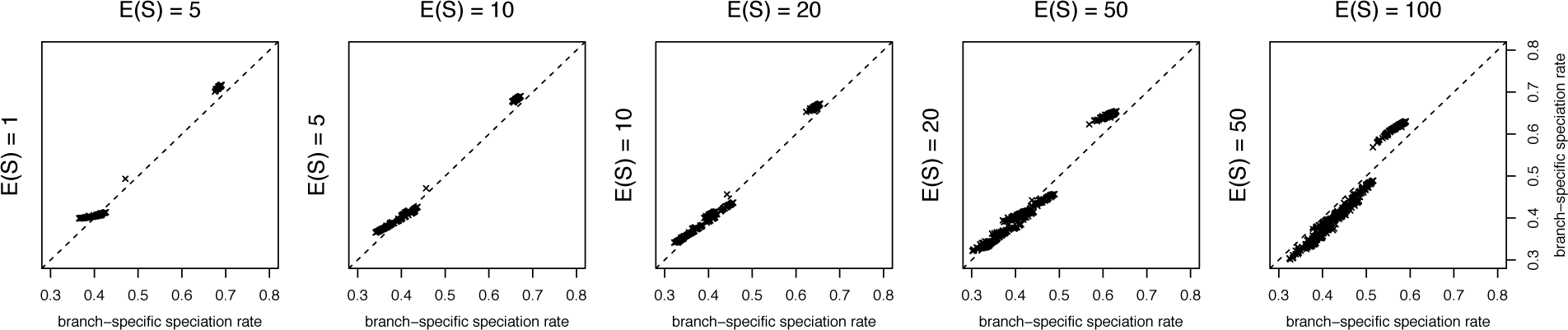
Comparison of branch-specific speciation-rate estimates between different priors on the expected number of diversification-rate shifts. We estimated the posterior mean speciation rate for each branch of the primate tree under different shift-rate priors, with the prior on *η* specified so that the prior expected number of rate-shift events under a Poisson process, *E*(*S*), was 1, 10, 20, 50 or 100. Despite the estimated number of diversification-rate shifts being prior sensitive (Figure 7), the branch-specific speciation-rate estimates are relatively robust to the prior on the expected number of diversification-rate shifts.

**Figure 14:**
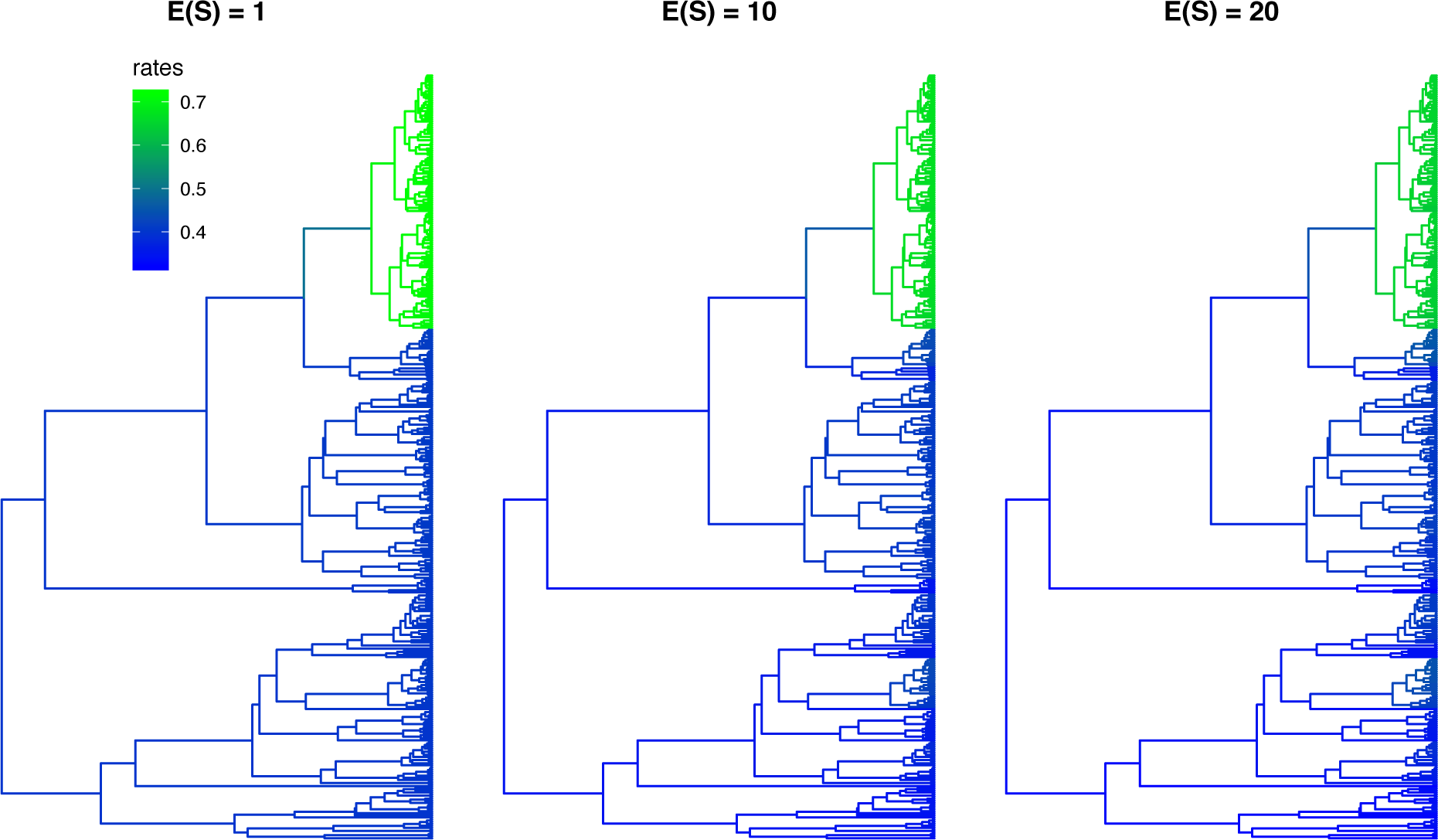
Branch-specific speciation-rate estimates for the primate tree under different shift-rate priors. We performed lineage-specific birth-death-shift analyses to estimate the posterior mean speciation rate for each branch of the primate tree under three different shift-rate priors, specified such that the expected number of diversification-rate shifts, *E*(*S*), was 1, 10, or 20. Branch colors reflect the branch-specific speciation-rate estimates; the scale bar is the same for all prior settings.

## Discussion

### Model Parameterization and Prior Specification

Our lineage-specific birth-death-shift process consists of three event types (speciation, extinction, and rate-shifts) which are governed by their respective rates. The speciation and extinction rates are drawn from some base distribution whereas the shift-rate is constant (*i.e.,* homogeneous) over the entire phylogeny. In this study we have taken a first step to explore the robustness of parameter estimates (*i.e.,* branch-specific diversification rates and the number of diversification-rate shifts).

In our analyses on simulated and empirical data we observed that the estimated number of diversification-rate shifts is sensitive to the choice of shift-rate prior (Figure 12). We have not explored the impact of the shape of the base distributions on the speciation and extinction rates. Instead, we emphasize that our implementation in RevBayes allows flexible parameterization of the lineage-specific birth-death-shift model. Here, we provide the foundation for further model exploration. We elaborate on the full flexibility of the model specification below.

#### Model parameterization

The lineage-specific birth-death-shift process defines a family of models that make different assumptions regarding the nature of diversification-rate variation across lineages. For example, the most general parameterization allows both speciation and extinction rates to vary independently across the tree. Under this model, a diversification-rate shift involves a change to new speciation and extinction rates that are independently drawn from their corresponding base distributions. From this model, two nested models can be specified: (1) a model that allows speciation rates to vary across the tree, but assumes a shared extinction rate for all branches (*i.e.,* diversification-rate shifts involve changes to the speciation rate), and (2) a second model that allows extinction rates to vary across the tree, but assumes a shared speciation rate for all branches (*i.e.,* diversification-rate shifts involve changes to the extinction rate). These models may also be parameterized using composite diversification-rate parameters, where diversification-rate shifts involve changes to the net-diversification rate, *r* = (*λ* − *µ*) and/or the relative-extinction rate, ϵ = (*µ* ÷ *λ*).

Finally, we could parametrize the lineage-specific birth-death-shift model where speciation and extinction rates are assumed to vary dependently across the tree. Under this model, a diversification-rate shift involves a change from one pair of rates (*λ*_*i*_,*µ*_*i*_), (where *i* corresponds to the same discrete rate category of both base distributions) to a new pair of speciation and extinction rates (*λ*_*j*_,*µ*_*j*_). For example, a diversification-rate shift might involve a change from paired rates (*λ*_3_,*µ*_3_) to (*λ*_5_, *µ*_5_) (reflecting a shift from the third to the fifth discrete categories of the speciation-and extinction-rate base probability distributions).

In RevBayes we provide full flexibility of applying any variant of how diversification rates change across lineages. It remains open to the biologist and further studies which type of diversification-rate variation is most prevalent and robust.

#### Prior distribution on the diversification rates

We adopt a Bayesian statistical approach to estimate the parameters of the lineage-specific birth-death-shift model. Therefore, we must specify a prior probability distribution for each parameter. Parameters of the lineage-specific birth-death-shift model are the speciation rate, *λ*, the extinction rate, *µ*, and the rate of diversification-rate shifts, *η*. Our implementation in RevBayes provides tremendous flexibility in the choice of priors for each parameter. For example, we might specify a lognormal, gamma, or exponential probability distribution as the prior on the speciation rate. Additionally, for a given choice of prior, RevBayes allows the user to either specify fixed values for the parameters of the chosen prior probability distribution (the ‘hyperparameters’), or to specify a more hierarchical Bayesian model by treating these hyperparameters as random variables (in which case we would specify a hyperprior for each hyperparameter). For example, if we chose a lognormal prior for the speciation rate, we could either specify fixed values for the parameters of this distribution (*i.e.,* the mean and standard deviation of the lognormal distribution), or we could specify hyperprior distributions for the mean and standard deviation hyperparameters. Thus, our implementation in RevBayes provides much more flexibility in specifying models compared with similar implementations (*e.g.,* BAMM only allows an exponential prior distribution with a fixed mean parameter for the speciation and extinction rate).

As a demonstration, we analyzed the primates phylogeny using a hierarchical model for the lognormal base distribution of the diversification rates. We assumed a uniform prior distribution between 0 and 100 for the mean of the lognormal base distribution and an exponential prior distribution with a mean of 0.587405 (we expect that 95% of the lognormal base distribution spans one order of magnitude; Höhna et al. 2017). Our example analysis shows that the hyperprior parameters of the base distribution can indeed be estimated (Figure 15). That is, the phylogeny appears to have sufficient information about the mean and variation branch-specific speciation rates. The hyperparameter estimates are not driven by their choices of prior distributions. Furthermore, the hierarchical approach reduces the prior sensitivity. Thus, we recommend to use such a hierarchical model for empirical analyses because it is difficult, if not impossible, to know which mean and standard deviation to assume for the base distribution of the diversification rates.

**Figure 15:**
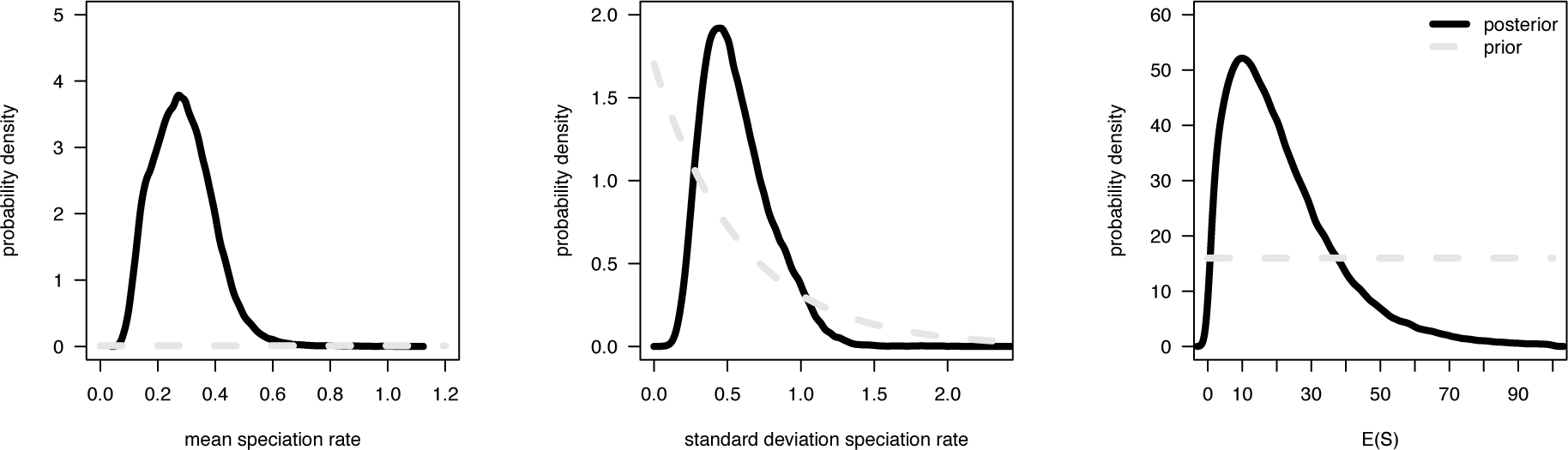
Estimation of hyperparameters under the hierarchical lineage-specific birth-death-shift model. We estimated the mean (*m*_*λ*_) and standard deviation (*sd*_*λ*_) of the lognormal base distribution for the speciation rate (left and middle panels). Additionally, we estimated the shift-rate *η* (right panel; showing the expected number of shifts, *E*(*S*), for an intuitive interpretation). We used the following prior distributions: *m*_*λ*_ ~ Unif(0, 100), *sd*_*λ*_ Exp(1.0/0.587405), and *E*(*S*) Unif(0, 100). The posterior distributions (black solid lines) show clear deviations from the corresponding prior distributions (light-gray dashed lines).

#### Prior Sensitivity and Estimating the Number of Rate Shifts

Our analyses have shown that the estimated number of diversification-rate shifts is very sensitive to the assumed prior distribution on the shift-rate (Figure 12). This prior sensitivity is actually expected because many small diversification-rate changes can have a similar effect as few large diversification-rate changes (Huelsenbeck et al. 2000). Our results do not imply that the shift-rate (and the number of diversification-rate shifts) is not identifiable. Specifically, Figure 12 shows that there is a (weak) signal for at least one diversification-rate shift but fewer than 20.

In practice, a biologist might have a good idea what number of diversification-rate shifts to expect for a given study group. However, we caution researchers to over-interpret the estimated number of diversification-rate shifts. We emphasize that in every empirical analysis either a set of prior assumptions should be applied (*e.g.,* by setting the number of *a priori* expected diversification-rate shifts to 1, 10 and 20), or a hyperprior distribution on the shift-rate *η* should be used. In our primate example analysis we observe that there is some signal for the shift-rate *η* (Figure 15; right panel). Moreover, the hyperprior analysis (Figure 15; right panel) confirms the results about the expected number of diversification-rate shifts of the fixed-prior analyses (Figure 12).

### Future directions and applications

In the present study we have focused on estimating branch-specific diversification rates. Nevertheless, our lineage-specific birth-death-shift model can be extended and applied in several different ways. Here we provide some thoughts to stimulate further ideas and research.

#### Correlation of diversification rates to other model components

We can extend the lineage-specific birth-death-shift model to analyses where other parts of the model (*e.g.,* rates of molecular or morphological evolution) are correlated with the speciation and extinction rates. Let us consider as an example an analysis where the rates of speciation correlate with the rate of molecular evolution by 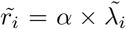 where 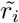 is the average rate of molecular evolution on branch *i* and 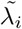 is the average rate of speciation on branch *i*. Then, we compute the average rate of speciation per branch 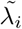 and deterministically transform these average speciation rates into average rates of molecular evolution 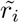. Thus, for this type of analyses we have to use the data-augmentation method because it adds the diversification-rate shifts onto the phylogeny. In such a situation, the rates of molecular evolution also have an impact on the number, location/timing and magnitude of the diversification-rate shifts. It is the joint posterior probability of the diversification-rate shifts, the speciation and extinction rates, and the rates of molecular evolution that we will estimate. Since the stochastic character mapping draws the diversification-rate shifts only from the lineage-specific birth-death-shift process without any information about other parts of the model depending on these events and rates, the stochastic character mapping method is not applicable in these types of analyses. However, the stochastic character mapping can be used as a proposal distribution in the MCMC algorithm. In RevBayes, such applications to linked models are readily available.

#### Cladogenetic and anagenetic diversification-rate shifts

The lineage-specific birth-death-shift process described here permits for shifts in speciation and extinction rates along the branches of a phylogeny (*i.e.,* anagenetic diversification rate shifts). However, many biological explanations for diversification-rate shifts have been hypothesized to correspond with speciation events (*i.e.,* cladogenetic diversification-rate shifts). Diversification-rate shifts have been modeled as occurring simultaneously with, for example, allopatric speciation events (Goldberg et al. 2011), cladogenetic changes in life history traits such as breeding system (Goldberg and Igić 2012), and cladogenetic changes in chromosome number or ploidy (Freyman and Höhna 2019). In contrast to those models, the lineage-specific birth-death-shift process tests for diversification rate shifts unassociated with an observed character. However, a biologist may want to use the lineage-specific birth-death-shift process to explore the possibility of diversification-rate shifts occurring at speciation events. Our stochastic character mapping approach for the lineage-specific birth-death-shift process is described primarily by Equations 3, 4, and 6, which represent a special case of the backward-and forward-time ODEs in Freyman and Höhna (2019) that enable both anagenetic and cladogenetic diversification-rate shifts. These general equations are implemented in RevBayes and could be used along with our approach discretizing the speciation-and extinction-rate base probability distributions into *k* categories to test a lineage-specific birth-death-shift process with both cladogenetic and anagenetic diversification-rate shifts. We mention this aspect of our implementation to highlight its flexibility for testing different diversification scenarios, however, we leave further exploration of cladogenetic diversification rate shifts to future work.

#### The lineage-specific birth-death-shift process as a prior distribution on divergence times

The primary goal of our development of the lineage-specific birth-death-shift process was to estimate branch-specific speciation and extinction rates. However, in RevBayes one can use the lineage-specific birth-death-shift process as a prior distribution on the phylogeny, *i.e.,* divergence times and tree topology. Recent studies have shown the impact of prior distributions on divergence times, although the overall importance is not fully understood (Condamine et al. 2015; Foster et al. 2016; Donoghue and Yang 2016). Allowing for rate variation among lineages is likely a more biologically realistic model and thus should be preferred. Using our lineage-specific birth-death-shift process in RevBayes it is now possible to estimate divergence time using this biologically more realistic model.

If the purpose of such an analysis is only to estimate the phylogeny and divergence times, then the lineage-specific birth-death-shift implementation integrating over all rate categories should be preferred (the stochastic character mapping step can be omitted). If instead the goal of the analysis is to estimate branch-specific speciation and extinction rates, as well as the phylogeny and divergence times, then the stochastic character mapping method should be used. The data-augmentation method has the fundamental problem that changes in the tree topology could consequently lead to changes in the the assignment of branches to rate categories. This problem also occurs if we would take phylogenetic uncertainty into account by using a sample of phylogenies from the posterior distribution (also available in RevBayes).

A major open issue is how to summarize branch-specific speciation and extinction rates for different phylogenies. Specifically, branches may have a different meaning for different phylogenies. More research is needed on how to interpret diversification-rate changes for different phylogenies.

## Conclusions

In the present paper we have introduced the lineage-specific birth-death-shift process, a stochastic branching process to model diversification rate variation among lineages. We described two different methods for estimating branch-specific speciation and extinction rates: data-augmentation and stochastic character mapping. We presented a validation of our implementation of the two methods in RevBayes and discussed potential applications and pitfalls. Most importantly, we provide researchers with a consistent model and correct implementation for estimating branch-specific speciation and extinction rates.

## Acknowledgements

This research was supported by the Deutsche Forschungsgemeinschaft (DFG) Emmy Noether-Program HO 6201/1-1 awarded to SH, and National Science Foundation (NSF) grants DEB-0842181, DEB-0919529, DBI-1356737, and DEB-1457835 awarded to BRM, and by DEB-1759909 awarded to JPH, and also by DEB-1655478 awarded to WAF.

